# Parkinson disease-associated protein DJ-1 regulates the autophagic-lysosomal pathway through ROS-dependent modulation of the AMPK/mTORC1 axis

**DOI:** 10.64898/2026.04.09.717467

**Authors:** Francesco Agostini, Federica De Lazzari, Alexandra Beilina, Bibiana Sgalletta, Chiara Sinisgalli, Elena Giusto, Laura Civiero, Luigi Bubacco, Mark R. Cookson, Nicoletta Plotegher, Elisa Greggio, Marco Bisaglia

## Abstract

Mutations in the protein DJ-1 are linked to familial forms of Parkinson’s disease (PD). The protein has been well-documented to exert a role in energy metabolism and antioxidant defense, contributing to the maintenance of mitochondrial homeostasis. We and others have previously observed that DJ-1 can also influence autophagy, but the mechanisms are still incompletely defined.

In this study, using complementary cellular and animal models, we characterize the impact of DJ-1 loss on the autophagic pathway. Our data demonstrate that DJ-1 deficiency impairs autophagosome-lysosome fusion and lysosomal degradation, resulting in the accumulation of dysfunctional autolysosomes and the subsequent buildup of autophagic substrates. Mechanistically, we show that elevated reactive oxygen species (ROS) in DJ-1-null models inhibit the energy-sensing AMP-activated protein kinase (AMPK), thereby activating the autophagy suppressor mechanistic target of rapamycin 1 (mTORC1). Collectively, these findings delineate a novel signaling axis linking oxidative stress to autophagic dysfunction, providing new insights into the cellular mechanisms underlying autophagic dysfunction in PD.

## Introduction

Parkinson disease (PD) is a neurodegenerative disorder that affects approximately 1% of the population over the age of 65 [1]. Approximately 5–10% of PD cases are monogenic [2], and analyses of familial forms of PD have provided important insights into the underlying pathogenic mechanisms. Proposed mechanisms include impairment of cellular clearance systems leading to protein misfolding and aggregation, mitochondrial dysfunction, and increased oxidative stress [1–3]. Several proteins associated with genetic forms of PD have been shown to exert direct or indirect roles in mitochondrial homeostasis [3,4]. Likewise, the involvement of lysosomes in PD pathology has gained strong support from genetic data [2–4].

In recent years, it has become increasingly evident that mitochondria and lysosomes are capable of reciprocal communication. Consequently, the activity of several PD-associated proteins is not restricted to a single organelle, and their functional impairment can affect both mitochondrial and lysosomal homeostasis. For example, although silencing of the lysosomal polyamine exporter ATP13A2 has been primarily associated with pronounced alterations in lysosomal function, impairment of this protein has also been linked to mitochondrial deficiencies, including decreased mitochondrial respiration [5]. Similarly, a decrease in glucocerebrosidase 1 (GCase) enzymatic activity, which is mainly associated with lysosomal dysfunction, also leads to compromised mitochondrial activity, including lower membrane potential, impaired mitochondrial respiration, and mitochondrial fragmentation [5]. Conversely, mutations in proteins associated with mitochondrial quality control have been described to also affect lysosomes. For instance, in addition to mitochondrial defects, PINK1 loss-of-function has been shown to induce inhibition of lysosomal activity and enlargement of lysosomal compartments [6], suggesting that this protein may influence the autophagic-lysosomal pathway (ALP) through indirect mechanisms. Moreover, patient-derived parkin-deficient fibroblasts exhibit a marked decrease in endosomal tubulation together with impaired retromer function [7].

The most direct form of communication between mitochondria and lysosomes is the physical interaction between these organelles [3,8]. Such contacts have been observed to participate in the regulation of several cellular processes, including mitochondrial fission and lysosomal dynamics [9]. In addition, these contact sites appear to be regions of intense metabolic activity, serving, for example, as hubs for the transfer of ions and small molecules between organelles [10]. Another form of direct communication is represented by mitophagy, the selective lysosomal degradation of mitochondria. Mitophagy depends on lysosomal activity and ensures maintenance of a functional mitochondrial pool through the removal of defective organelles [3,8]. Beyond direct organelle interactions, it has also been demonstrated that long-distance communication between mitochondria and the ALP exists and can be mediated by specific signaling cascades. Among them, phosphorylated (active) AMPK [11], one of the principal sensors of intracellular energetic status and autophagy regulator [12], is reduced in a mitochondrial dysfunction cell model based on stable knockdown of a respiratory chain complex III subunit. Alternatively, mTORC1-dependent inhibition of the MiT family of transcription factors, which in turn promotes lysosomal biogenesis and autophagy, has been proposed as a key point of convergence between mitochondrial function and the ALP [8].

Despite the growing attention devoted to mitochondria–lysosome crosstalk in PD pathology, the mechanisms underlying long-distance communication between these organelles remain largely unexplored. Among the proteins associated with genetic forms of PD that may be exploited to investigate such mechanisms, DJ-1 represents a particularly promising candidate. Long-standing evidence supports the involvement of DJ-1 in maintaining mitochondrial homeostasis. Indeed, DJ-1 has been shown to sustain mitochondrial respiration and prevent excessive mitochondrial-derived reactive oxygen species (ROS) production, and its absence has been shown to induce mitochondrial fragmentation and a reduction in mitochondrial membrane potential [13,14]. Furthermore, DJ-1 has been proposed to directly modulate mitochondrial activity by sustaining the function of respiratory chain complexes I and V [15–17]. In addition, DJ-1 has been implicated in the regulation of mitochondrial dynamics and in the control of mitophagy [13,14,18–24]. Most mitochondrial phenotypes associated with DJ-1 loss-of-function might be explained by a recently identified physiological role of this protein. Compelling evidence has demonstrated that DJ-1 can catalyze the hydrolysis of highly reactive cyclic 3-phosphoglyceric anhydride, a metabolite of glycolysis, capable of reacting with and modifying cytosolic molecules, including glutamate, glutathione, and lysine and cysteine residues in proteins [25–27]. Although the proportion of modified versus unmodified targets has not yet been quantified, the progressive accumulation of modified proteins in post-mitotic neuronal cells is likely to impair proteostasis, thereby promoting ROS accumulation and mitochondrial dysfunction.

Beyond its role in mitochondrial homeostasis, several studies also suggest that DJ-1 affects the ALP, although reports describing its function at the autophagic level are often inconsistent. Experimental evidence supports a correlation between DJ-1 deficiency and reduced autophagy [18,28,29], whereas other studies have reported opposite effects [15,30]. These discrepancies may derive from the fact that autophagy is a highly complex and dynamic process, and monitoring LC3 lipidation alone, as often reported, may be insufficient to unambiguously assess the direction of autophagic flux. For this reason, additional experimental parameters must be considered to clarify whether and how DJ-1 regulates the autophagic pathway. Recently, using a *Drosophila melanogaster* model of familial PD based on DJ-1 knockout, we demonstrated that loss of the fly DJ-1 ortholog leads to hypoactivity and increased susceptibility to food deprivation, supporting the presence of metabolic impairments and suggesting a role for DJ-1 as a regulator of energy balance at the crossroads between mitochondrial and autophagic pathways [31]. DJ-1 indeed appeared to contribute to mitochondrial homeostasis by sustaining organelle morphology and complex I respiration, while simultaneously participating in autophagy regulation [31].

In the present work, we employed multiple DJ-1 loss-of-function experimental models to characterize the role of DJ-1 in the autophagic–lysosomal pathway. In these models, we also investigated the involvement of AMPK and mTORC1, as the signaling pathways that may mediate the crosstalk between mitochondria and lysosomes, defining alterations in the intracellular redox state as one of the mechanisms associated with their modulation.

## Results

### The absence of DJ-1 affects autophagy in mouse embryonic fibroblasts (MEFs)

Cellular degradation processes allow the removal of cellular debris, such as aged or damaged organelles and dysfunctional or misfolded proteins. By analyzing the levels of degradative substrates, it is possible to obtain an indication of the clearance activity of a biological system. Ubiquitin is a small protein that can covalently bind to other proteins and is fundamental in promoting their autophagy-dependent degradation [32]. Consequently, the cellular levels of ubiquitin and ubiquitylated proteins vary according to their degradation rate. Therefore, in the first series of experiments, we evaluated the levels of these clearance substrates in DJ-1 knockout (KO) MEFs. As shown in Fig. 1A, DJ-1 KO cells are characterized by higher levels of ubiquitylated proteins compared to control cells, suggesting an impairment of clearance processes upon DJ-1 deletion. We then assessed the levels of the lipidated form of LC3 (microtubule-associated protein 1A/1B-light chain 3), a key autophagosomal marker involved in cargo recognition [33], as well as those of the autophagy adaptor protein p62, which binds both LC3 and ubiquitylated cargoes to ensure their delivery to degradation [34]. Higher levels of lipidated LC3 were detected in DJ-1 KO cells relative to wild-type (WT) cells (Fig. 1B), suggesting an enhancement in autophagosome number, possibly due to alterations in autophagosomal generation and/or turnover. In contrast, DJ-1 deletion did not significantly affect p62 levels.

**Figure 1.**
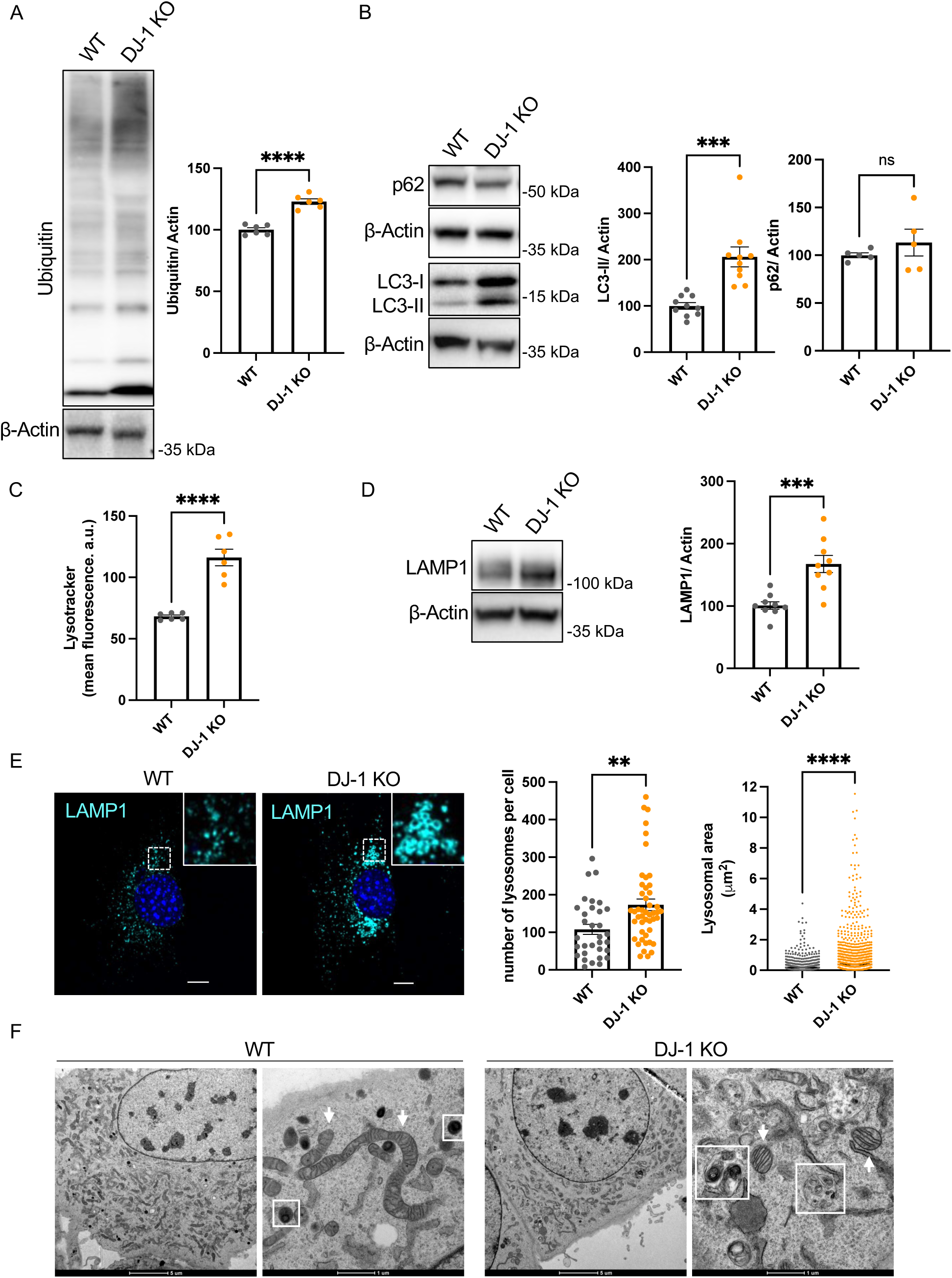
The absence of DJ-1 affects the autophagy-lysosomal pathway in MEF cells. (A) Western blot analysis of ubiquitinated proteins in WT and DJ-1 KO MEF. Six independent biological replicates were analyzed. Statistical significance was determined using an unpaired t-test (t: 8.324; df:10; ****p < 0.0001). (B) Western blot analysis of p62 and LC3 in WT and DJ-1 KO MEF. At least 5 biological replicates were analyzed. Statistical differences were evaluated by an unpaired t-test (p62: t test with Welch ‘s correction, t: 0.9278; df: 4.269; p: not significant; lipidated LC3 (LC3-II): t: 4.705; df: 18; *** p < 0.001). (C) Quantification of mean LysoTracker-green fluorescence intensity by flow cytometry. Six independent biological replicates were analyzed. Statistical significance was determined by an unpaired t-test (**** p < 0.0001). (D) Western blot analysis of LAMP1 in WT and DJ-1 KO MEF. 9 biological replicates were analyzed. Statistical significance was determined using an unpaired t-test (t: 4.433; df: 16; *** p < 0.001). (E) Confocal imaging of LAMP1 in WT and DJ-1 KO MEF. Scale bar: 10 μm. The graphs on the right show the quantification of lysosome number per cell (at least 30 cells across three independent replicates, unpaired t-test, t: 3.020; df: 77; ** p < 0.01) and lysosomal size (more than 1000 lysosomes analyzed; unpaired t-test; t:8.478; df: 4533; **** p < 0.0001) (F) TEM images of WT and DJ-1 KO MEF. Scale bar: 5 μm (low magnification) and 1 μm (high magnification). The white boxes indicate autophagic vesicles in WT cells and autophagic vesicles containing undigested materials in DJ-1 KO cells. White arrows indicate mitochondria.

Considering the observed variation in autophagosome abundance, we next investigated whether lysosomes were also affected. Cells were first stained with LysoTracker, a cell-permeable fluorescent dye that labels acidic intracellular compartments, and fluorescence was subsequently quantified by flow cytometry. DJ-1 KO cells exhibited a higher LysoTracker signal than WT cells, consistent with an enhanced number of lysosomes and/or higher lysosomal size (Fig. 1C). To further validate the increase in the lysosomal compartment in DJ-1 KO cells, we analyzed the levels of LAMP1 (lysosomal-associated membrane protein 1), a glycosylated protein essential for lysosomal membrane integrity and function [35]. Consistent with LysoTracker staining, higher LAMP1 levels were detected by western blot (WB) in DJ-1 KO cells compared to control cells (Fig. 1D). Immunohistochemical analyses further revealed a greater number of LAMP1-positive structures, many of which appeared larger than those observed in control cells (Fig. 1E). These observations were further supported by transmission electron microscopy. In line with previous reports [18], DJ-1 KO cells displayed structures containing electron-dense material, which were absent in control cells (Fig. 1F) and have been previously described as undigested cellular components [18]. Notably, DJ-1 KO cells also exhibited alterations in mitochondrial morphology, characterized by shorter and more rounded mitochondria, consistent with previous cellular and *in vivo* findings [13,14,31].

### Lysosomal activity and autophagic flux are impaired in DJ-1 KO MEFs

Since lysosomes represent crucial players in the intracellular catabolic pathways, dysfunctions in their activity can lead to clearance defects and accumulation of autophagic substrates. Thus, we next evaluated lysosomal degradative capacity using two independent assays. First, we performed the MagicRed assay, which enables quantification of cathepsin B activity. The MagicRed probe is a non-toxic lysosomal substrate that becomes fluorescent upon cleavage by active cathepsin B enzyme. Consistent with the LAMP1-associated signal, larger structures were observed in DJ-1 KO cells, compatible with the presence of enlarged lysosomal structures. Mean MagicRed fluorescence intensity was normalized to total MagicRed-positive area, revealing lower levels of proteolytic activity in the absence of DJ-1 compared to WT cells (Fig. 2A). To independently confirm DJ-1-associated lysosomal dysfunction, we conducted DQ-BSA assays, which utilize a protease substrate that emits fluorescence upon enzymatic cleavage. The fluorescence intensity measured by flow cytometry directly correlates with degradative activity [36]. Given the previously observed lysosomal accumulation, cells were co-treated with DQ-BSA and LysoTracker, and DQ-BSA-derived fluorescence was normalized to the LysoTracker fluorescence intensity. In agreement with the MagicRed assay, DJ-1 KO cells displayed lower lysosomal protease activity per lysosomal unit compared to control cells, indicating impaired lysosomal degradative capacity due to DJ-1 loss (Fig. 2B).

**Figure 2.**
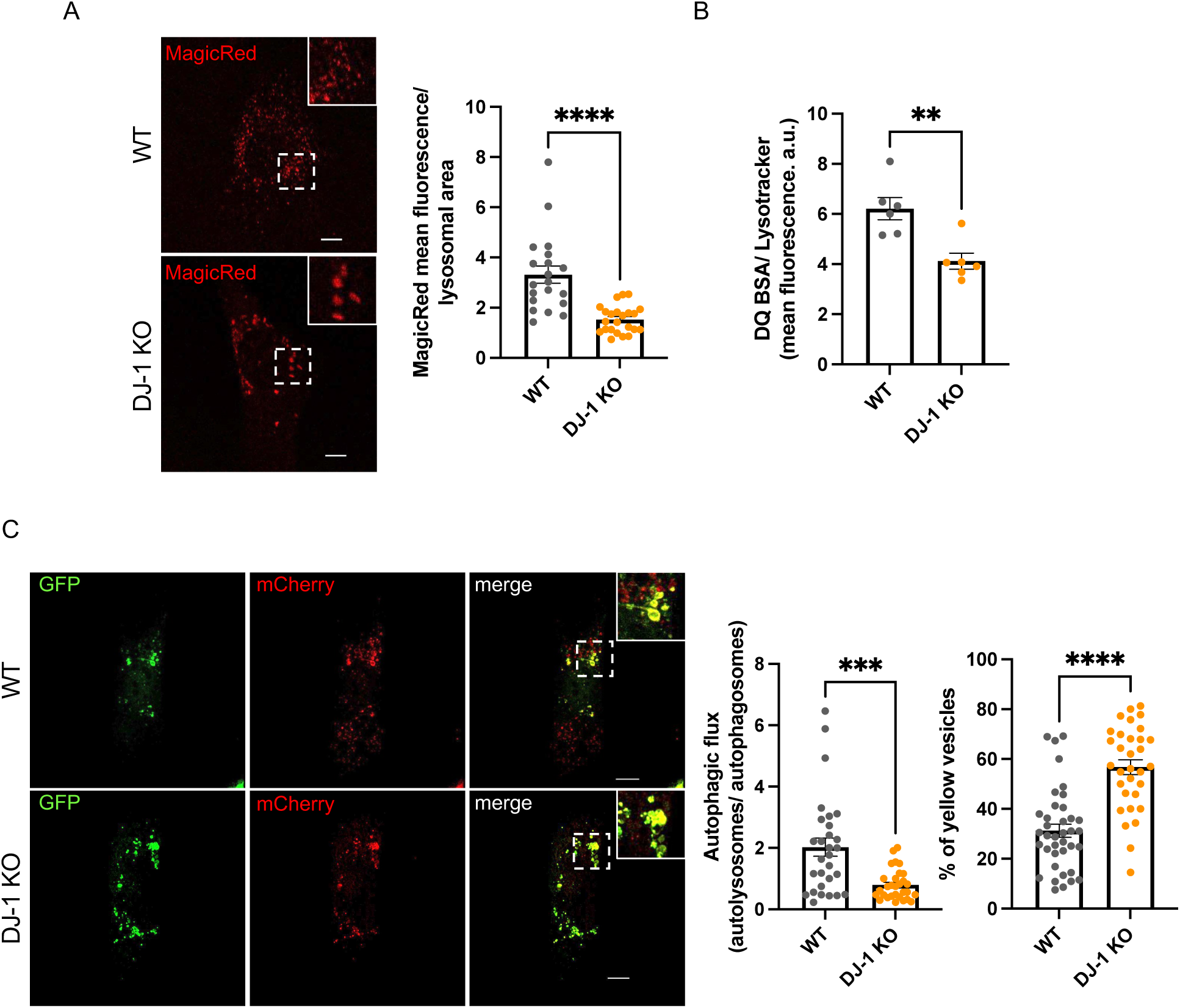
The absence of DJ-1 alters lysosomal activity and autophagic flux. (A) Live confocal imaging of WT and DJ-1 KO MEF exposed to the cathepsin B activity reporter MagicRed. Scale bar: 10 μm. The right graph shows quantification of MagicRed mean fluorescence normalized to MagicRed-positive area. At least 20 cells per genotype across 3 independent experiments were analyzed. Statistical differences were determined by an unpaired t-test (t: 5.185; df: 41; **** p < 0.0001). (B) Flow cytometric analysis of WT and DJ-1 KO MEF treated with the lysosomal protease activity reporter DQ BSA. Mean fluorescence intensity of DQ BSA was normalized to lysosomal content measured by LysoTracker-green fluorescence. Six independent biological replicates were analyzed using an unpaired t-test (t: 3.847; df: 10; ** p < 0.01). (C) Live confocal imaging of WT and DJ-1 KO MEF expressing the autophagic flux reporter mCherry-GFP-LC3. Scale bar: 10 μm. More than 25 cells per genotype across 3 independent experiments were analyzed. Panels on the right show quantification of the autophagic flux expressed as the ratio between red-only vesicles and yellow vesicles, and the percentage of autophagosomes (yellow vesicles). Statistical significance was determined using an unpaired t-test (autophagic flux: t: 4.018; df: 57; *** p < 0.001; percentage of autophagosomes: t: 6.430; df: 68; **** p < 0.0001).

To further assess whether autophagic-lysosomal flux was altered in the absence of DJ-1, we overexpressed the mCherry-LC3-GFP reporter in both control and DJ-1 KO cells. This construct consists of the autophagosomal membrane protein LC3 tagged with both GFP and mCherry, allowing independent visualization of autophagosomes and autolysosomes [37]. When LC3 localizes to autophagosomal membranes, GFP and mCherry fluorescence overlap, generating a yellow signal. Upon fusion with lysosomes, the acidic pH of autolysosomes quenches GFP fluorescence, leaving only the mCherry red signal. The ratio of red-only puncta (autolysosomes) to yellow puncta (autophagosomes) thus reflects autophagic flux. As shown in Fig. 2C, WT cells displayed a higher autophagic flux than DJ-1 KO cells. In contrast, DJ-1 KO cells exhibited a higher percentage of yellow puncta compared to WT cells, indicating autophagosome accumulation, likely due to impaired autophagosome–lysosome fusion. Collectively, these data suggest a defect in autophagosome–lysosome fusion and the accumulation of autolysosomal structures with lower degradative capacity, ultimately leading to autophagy substrate accumulation in basal conditions.

### The absence of DJ-1 alters the AMPK and mTORC1 axis in MEFs

Two pathways are primarily involved in modulating autophagy. The first one relies on the protein kinase mTORC1, whose activity inhibits autophagy [38]. The second mechanism depends on the kinase AMPK that can activate autophagy both indirectly, by inhibiting mTORC1, and directly by promoting autophagy initiation [39,40]. Thus, we verified whether DJ-1 can affect these two pathways. First, we assessed the activity of AMPK by evaluating the level of the phosphorylated form of the protein at threonine 172 (Thr172), which is well known to correlate with protein activity [40]. As represented in Fig. 3A, DJ-1 KO cells are characterized by lower levels of phosphorylated AMPK (p-AMPK) in comparison to WT cells, indicating an impairment in the cellular pathway regulated by this kinase. Then, to indirectly assess mTORC1 activity, we analyzed the phosphorylation of ribosomal protein S6 kinase beta-1 (S6K) at Thr389, a well-established mTORC1 target involved in protein synthesis and cell growth [41]. WB analyses revealed higher levels of phosphorylated S6K (p-S6K) in DJ-1 KO cells compared to controls (Fig. 3B), indicating enhanced mTORC1 pathway activation.

**Figure 3.**
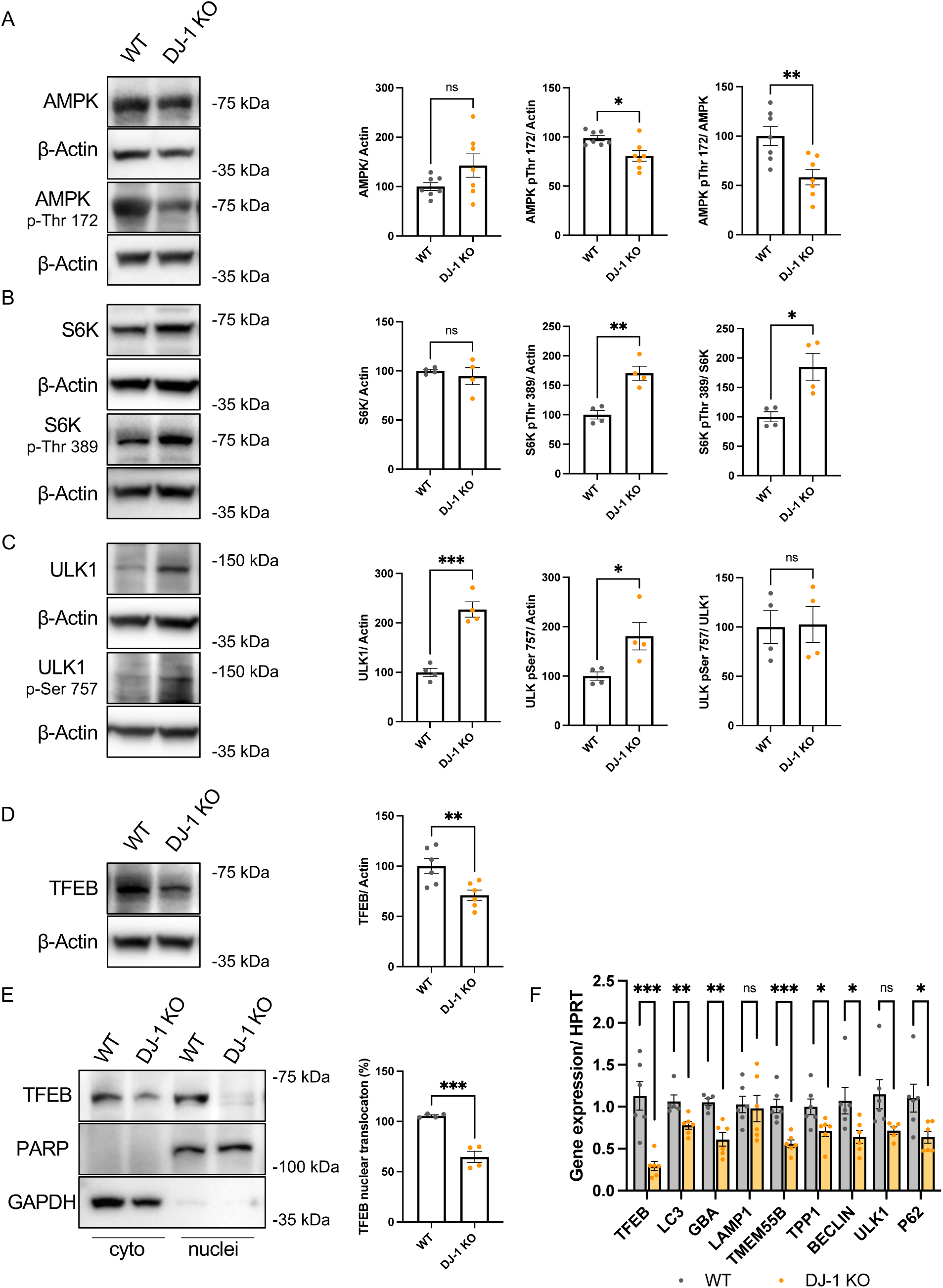
DJ-1 regulates the mTORC1-TFEB signaling axis. (A) Western blot analysis of AMPK total and phosphorylated in Thr172 (protein activation site) in WT and DJ-1 KO MEF cells. Seven biological replicates were analyzed. The graphs on the right show the quantification of total AMPK (normalized to the housekeeping protein actin) and phosphorylated AMPK (normalized to actin and to total AMPK level). Statistical significance was determined by an unpaired t-test (total AMPK: t: 1.708; df: 12; not significant; p-AMPK (normalized to actin): t: 3.030; df: 12; ** p<0,01; p-AMPK (normalized to total AMPK): t: 3.375; df: 12; ** p < 0.01). (B) Western blot analysis of total and phosphorylated S6K (Thr389, mTORC1 target site), downstream target of mTORC1, in WT and DJ-1 KO MEF. Four independent biological replicates were analyzed. The graphs show the quantification of total S6K (normalized to the housekeeping protein actin) and phosphorylated S6K (normalized to actin and to total S6K level). Statistical significance was determined by an unpaired t-test (total S6K: t: 0.6052; df: 6; not significant; p-S6K (normalized to actin): t: 5.048; df: 6; ** p<0,01; p-S6K (normalized to total S6K): t: 3.495; df: 6; * p < 0.05). (C) Western blot analysis of total and phosphorylated ULK1 (Ser757, mTORC1 target site) in WT and DJ-1 KO MEF. Four independent replicates were analyzed. Quantification of total ULK1 (normalized to actin), phosphorylated ULK1 (normalized to actin), and phosphorylated ULK1 (normalized to total ULK1) is shown below. Statistical significance was determined using an unpaired t-test (total ULK1: t: 4.161; df: 6; *** p < 0.001; p-ULK1 (normalized to actin): t: 3.643; df: 6; * p < 0.05; p-ULK1 (normalized to total ULK1): t: 1.391; df: 6; not significant). (D) Western blot analysis of TFEB protein levels in WT and DJ-1 KO MEF. Six biological replicates were analyzed. Statistical significance was determined by an unpaired t-test (t: 3.204; df: 10; ** p < 0.01). (E) Western blot analysis of TFEB levels in cytoplasmic and nuclear fractionations from WT and DJ-1 KO MEF. Cytoplasmic TFEB was normalized to the housekeeping GAPDH, while nuclear TFEB was normalized to the nuclear protein PARP. TFEB nuclear translocation was calculated as the ratio of nuclear TFEB to total TFEB. Four independent samples were used for the analysis. Statistical significance was determined using an unpaired t-test (t: 7.227; df: 6; *** p < 0.001). (F) Real-time PCR analysis of TFEB downstream genes in WT and DJ-1 KO MEF. Gene expression levels were normalized to the housekeeping gene HPRT. At least 5 biological replicates were analyzed. For each gene, the statistical significance between WT and DJ-1 KO cells were determined by unpaired t-test (TFEB: t: 4.679; df: 10; *** p < 0.001; LC3: t: 3.361; df: 9; ** p < 0.01; GBA: t: 4.610; df: 9; ** p < 0.01; LAMP1: t: 0.2568; df: 10; not significant; TMEM58B: t: 4.781; df: 10; *** p < 0.001; TPP1: t: 2.517; df: 10; * p < 0.05; Beclin1: t: 2.459; df: 10; * p < 0.05; ULK1: t: 2.215; df: 9; not significant; p62: t: 2.578; df: 10; * p < 0.05).

To further characterize the involvement of mTORC1 signaling in DJ-1-dependent ALP impairment, we examined two established downstream autophagic effectors of mTORC1, the kinase ULK1 and the transcription factor TFEB, both of which are inhibited by mTORC1-mediated phosphorylation. Upon autophagy induction, unphosphorylated ULK1 regulates early steps of autophagosome formation. In DJ-1 KO cells, we observed higher levels of inactive phospho-ULK1, further supporting elevated mTORC1 activity (Fig. 3C).

TFEB is also phosphorylated by mTORC1, leading to its cytoplasmic retention via interaction with 14-3-3 proteins. This interaction inhibits nuclear translocation and promotes TFEB degradation through the ubiquitin–proteasome system, mediated by the chaperone-dependent E3 ubiquitin ligase STUB1 [42]. As shown in Fig. 3D, DJ-1 KO cells exhibited a marked reduction in total TFEB levels compared to controls. Moreover, consistent with enhanced mTORC1 activity, the fractionation of cytoplasmic and nuclear compartments in MEF cells also revealed a significant decrease in the nuclear translocation of TFEB, measured as the ratio of nuclear TFEB over total TFEB (Fig. 3E). To evaluate the functional consequences of this effect, we measured the expression of several TFEB-regulated genes, including TFEB itself [43]. Consistent with lower TFEB activity, most analyzed transcripts were downregulated in the absence of DJ-1 (Fig. 3F). Overall, these findings indicate that DJ-1 loss leads to a lower activation of AMPK and enhanced mTORC1 activation, resulting in TFEB-impaired nuclear translocation, and increased degradation, ultimately causing downregulation of autophagic-lysosomal genes and ALP dysfunction.

To determine whether the effects described above were specifically due to the absence of DJ-1, we reintroduced the protein into DJ-1 KO MEF and assessed the activation of the mTORC1 and AMPK pathways by WB. Notably, re-expression of DJ-1 in KO MEF restored AMPK activity (Fig. 4A) and reduced mTORC1 signaling (Fig. 4B) to levels comparable to those observed in control cells. Consistently, DJ-1 re-expression also normalized the levels of the lysosomal marker LAMP1, similar to those detected in WT cells (Fig. 4C).

**Figure 4.**
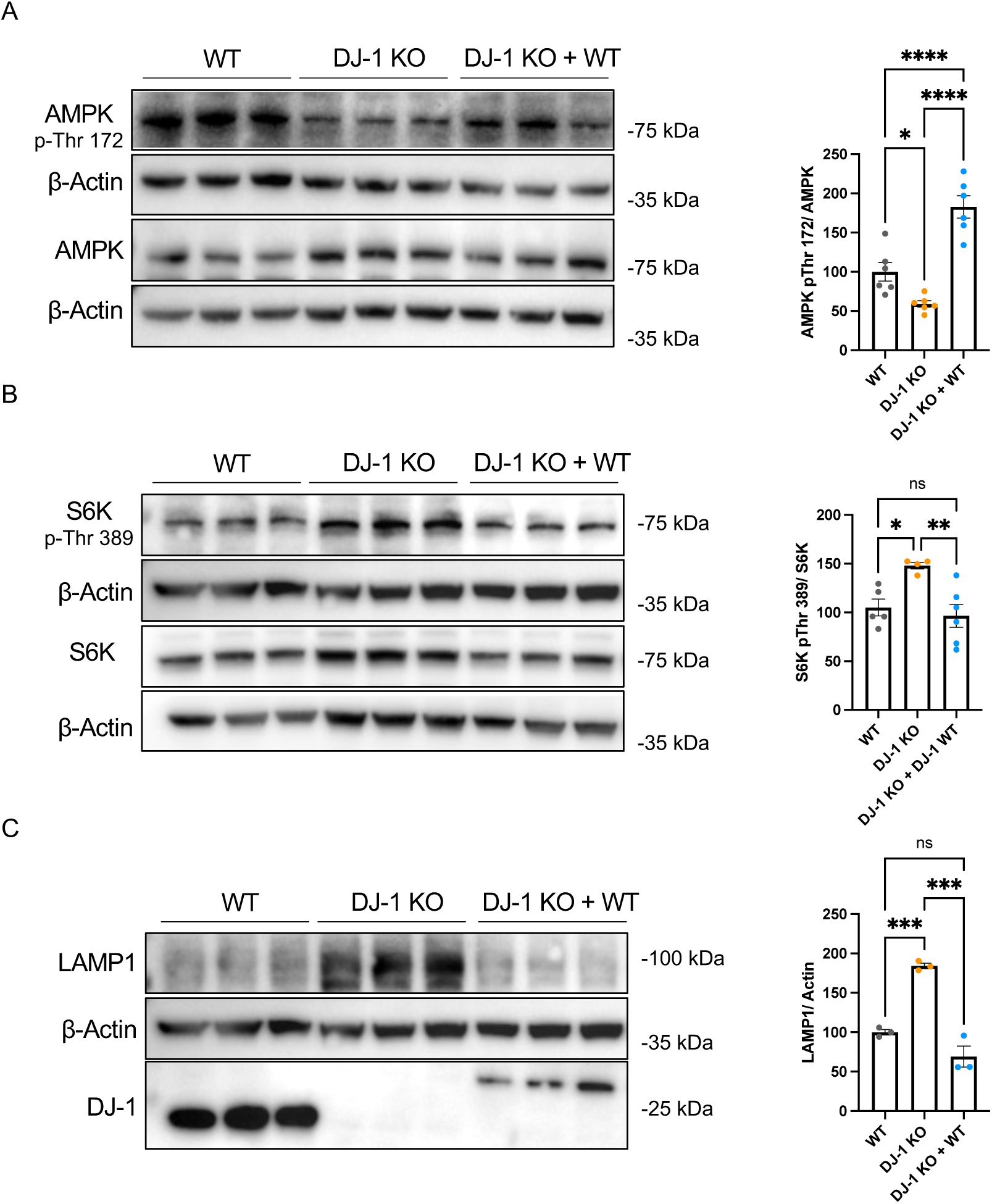
exogenous DJ-1 expression in DJ-1 KO cells rescues the phenotypes associated with the absence of the protein. (A) Western blot of total AMPK and AMPK phosphorylated in Thr172 in MEF cells WT, DJ-1 KO and DJ-1 KO transiently expressing DJ-1 WT. The graph on the right shows the level of phosphorylated AMPK normalized with the total level of the protein. Six independent samples were used for the analysis. Statistical significance was determined using a one-way ANOVA with Tukey’s multiple comparisons test (WT vs DJ-1 KO: df: 15; * p < 0.05; WT vs DJ-1 KO + DJ-1 WT: df: 15; *** p < 0.001; DJ-1 KO vs DJ-1 KO + DJ-1 WT: df: 15; **** p < 0.0001) (B) Western blot of total S6K and SK6 phosphorylated in Thr389 in MEF cells WT, DJ-1 KO and DJ-1 KO transiently expressing DJ-1 WT. The graph on the right shows the level of phosphorylated S6K normalized with the total level of the protein. At least 5 independent samples were used for the analysis. Statistical significance was determined using a one-way ANOVA with Tukey’s multiple comparisons test (WT vs DJ-1 KO: df: 12; * p < 0.05; WT vs DJ-1 KO + DJ-1 WT: df: 12; not significant; DJ-1 KO vs DJ-1 KO + DJ-1 WT: df: 12; ** p < 0.01). (C) Western blot of LAMP1 and DJ-1 in MEF cells WT, DJ-1 KO and DJ-1 KO transiently expressing DJ-1 WT. exogenous DJ-1 migrates at higher molecular weight in SDS page because it is tagged with V5-6xHis. The graph on the right shows the level of LAMP 1. Three independent biological replicates were included in the analysis. Statistical significance was determined using a one-way ANOVA with Tukey’s multiple comparisons test (WT vs DJ-1 KO: df: 6; *** p < 0.001; WT vs DJ-1 KO + DJ-1 WT: df: 6not significant; DJ-1 KO vs DJ-1 KO + DJ-1 WT: df: 6; *** p < 0.001).

### DJ-1 loss-of-function induces autophagic-lysosomal alterations in induced pluripotent stem cells differentiated into dopaminergic neurons, affecting both APMK and mTORC1 pathways

To validate DJ-1-dependent effects in a human-relevant model, we investigated autophagic alterations in dopaminergic neurons derived from induced pluripotent stem cells (iPSCs). We employed both WT human cells and an isogenic line carrying the p.Ala111LeuFs*7 DJ-1 mutation [44], which results in the complete absence of the protein, as shown in Fig. 5A. Due to their capacity to differentiate into neuronal lineages, iPSCs represent a highly relevant model to study DJ-1-associated mechanisms in a neuronal context. Accordingly, we performed WB analyses to assess autophagic markers in iPSCs differentiated into early midbrain dopaminergic neurons. Successful differentiation was confirmed by detection of the dopaminergic marker tyrosine hydroxylase (TH) (Fig. 5A). Consistent with MEF data, we detected an increase in ubiquitin level in DJ-1 KO TH-positive dopaminergic neurons (Fig. 5B) compared to control cells. Moreover, DJ-1 loss-of-function iPSC-derived neurons displayed higher levels of lipidated LC3 than control cells, while, as in MEF, p62 levels were not significantly altered (Fig. 5C). Similarly, DJ-1-deficient iPSC-derived neurons showed elevated levels of the lysosomal protein LAMP1 compared to control cells (Fig. 5D). Confocal microscopy analyses confirmed a significant increase in LAMP1 and LC3 mean fluorescence intensity in DJ-1 KO TH-positive cells (Fig. 5E). Lysosomal activity was then assessed using the DQ-BSA assay. In agreement with MEF results, DJ-1 loss affected the lysosomal protease activity in iPSC-derived neurons, confirming an impaired lysosomal degradative capacity (Fig. 5F).

**Figure 5.**
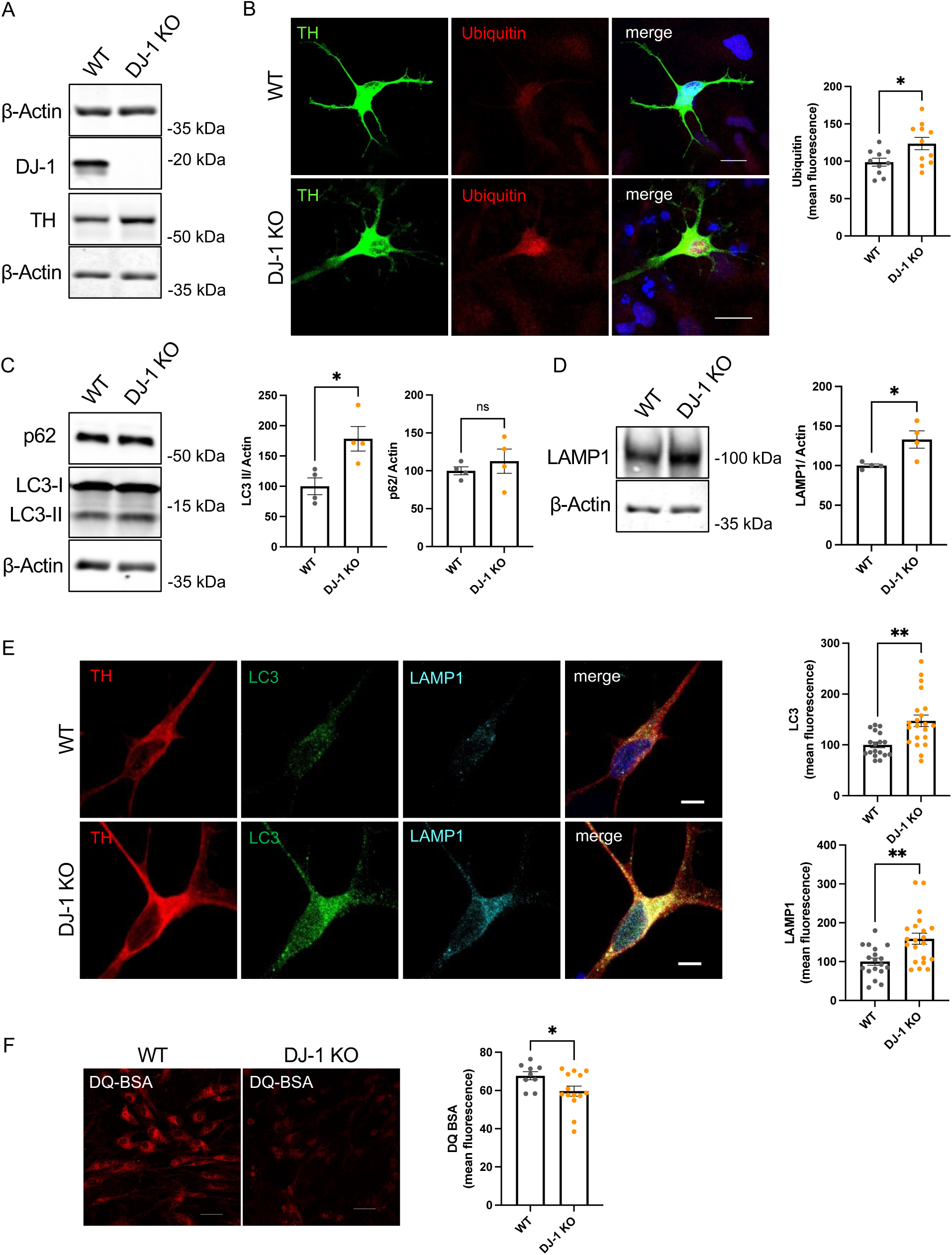
DJ-1 regulates autophagy and lysosomal activity in iPSC-derived dopaminergic neurons. (A) Western blot of DJ-1 and TH in WT and DJ-1 KO iPSC-derived neurons after differentiation into dopaminergic neurons. (B) Confocal images of WT and DJ-1 KO dopaminergic neurons stained for TH and ubiquitin. Scale bar: 10 μm. The mean ubiquitin fluorescence within TH-positive cells was quantified. At least 10 cells derived from 2 independent dopaminergic neuron differentiation experiments were used. Statistical significance was determined using an unpaired t-test (t: 2.474; df: 19; * p < 0.05). (C) Western blot analysis of p62 and LC3 in WT and DJ1 KO iPSC-derived neurons. Four biological samples derived from 2 independent dopaminergic neuron differentiation experiments were analyzed. Quantification of p62 and lipidated LC3 (LC3 II) levels is shown in the panels. An unpaired t-test was used for the statistical analysis (p62: t: 0.7589; df: 6; not significant; LC3 II: t: 3.200; df: 6; * p < 0.05). (D) Western blot analysis of LAMP1 in WT and DJ-1 KO iPSC-derived neurons. The analysis was performed using 4 biological samples derived from 2 independent dopaminergic neuron differentiation experiments. Statistical significance was evaluated by an unpaired t-test (t: 2.967; df: 6; * p < 0.05). (E) Confocal imaging of dopaminergic neurons stained for TH, LC3 and LAMP1. Scale bar: 10 μm. At least 18 cells from 2 independent dopaminergic neuron differentiation experiments were included in the analysis. The graphs on the right show the quantification of the LAMP1 and LC3 mean fluorescence intensity within TH-positive cells. Statistical analysis was performed using an unpaired t-test (LC3: t: 3.561; df: 37; ** p < 0.01; LAMP1: t: 3.365; df: 37; ** p < 0.01). (F) Live confocal imaging of WT and DJ-1 KO iPSC-derived neurons treated with the lysosomal protease activity reporter DQ-BSA. Scale bar: 50 μm. The mean DQ-BSA fluorescence intensity was quantified in every confocal field. The analysis includes at least 10 fields of cells derived from 2 independent dopaminergic neuron differentiation experiments. Statistical analysis was performed using an unpaired t-test (t: 2.167; df: 21; * p < 0.05).

We then assessed whether the DJ-1 loss-of-function affects AMPK and mTORC1 activity. Interestingly, also in dopaminergic neurons, the levels of p-AMPK, as evaluated by immunofluorescence analysis, were lower in DJ-1 KO cells compared to control (Fig. 6A). The activity of mTORC1 was then evaluated by measuring p-S6K levels, and, as observed in MEFs, DJ-1 loss-of-function iPSC-derived neurons exhibited a trend toward higher p-S6K levels than WT cells (Fig. 6B),

**Figure 6.**
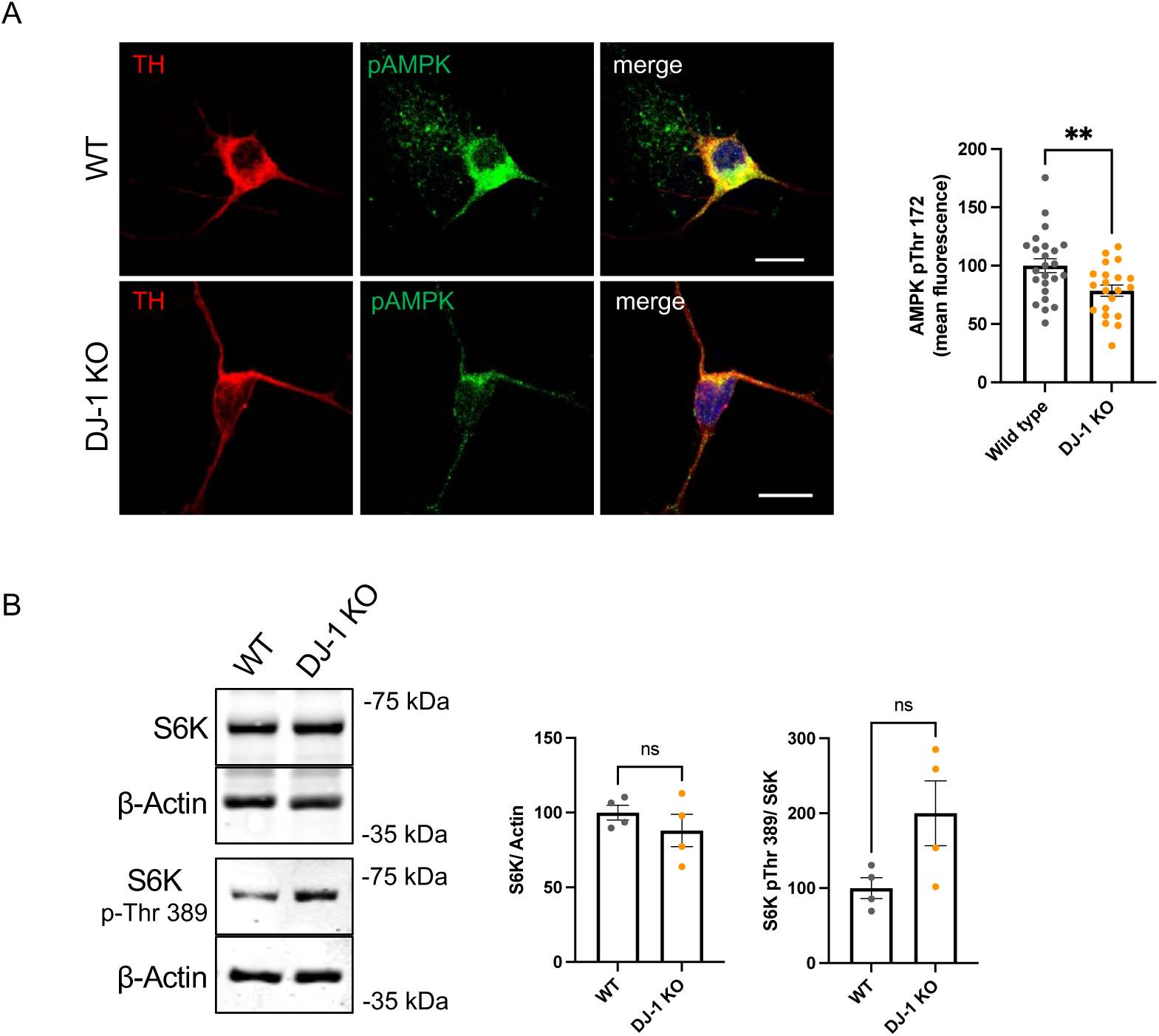
DJ-1 modulates the AMPK-mTORC1 pathway in iPSC-derived dopaminergic neurons. (A) Confocal imaging of dopaminergic neurons stained for the dopaminergic marker TH and phosphorylated AMPK. Scale bar: 10 μm. At least 21 cells deriving from 2 independent dopaminergic neuron differentiation experiments were included in the analysis. The graph shows the quantification of the phospho-AMPK mean fluorescence intensity within TH-positive cells. Statistical analysis was performed using an unpaired t-test (t: 2.775; df: 43; ** p < 0.01). (B) Western blot analysis of total and phosphorylated S6K (Thr389, mTORC1 target site) in WT and DJ-1 KO iPSC-derived neurons. Four biological samples derived from 2 independent dopaminergic neuron differentiation experiments were used. The graphs on the right show the quantification of total S6K (normalized to actin) and phosphorylated S6K (normalized to total S6K). Statistical significance was determined through an unpaired t-test (S6K: t: 1.006; df: 6; not significant; p-S6K: t: 2.202; df: 6; not significant).

### The absence of the DJ-1 ortholog affects the autophagic-lysosomal pathway in fruit flies

To validate our findings in an *in vivo* model, we employed *Drosophila melanogaster*. The fly genome encodes two DJ-1 homologs, dj-1α and dj-1β. While dj-1α expression is largely restricted to testes, previous studies have demonstrated that dj-1β is ubiquitously expressed and confers protection against oxidative stress, suggesting that it more closely recapitulates human DJ-1 function [45,46]. Accordingly, we utilized the dj-1β^Δ93^ strain, a null mutant carrying a 1960 bp deletion in the dj-1β gene, which was previously described and validated [45]. The w^1118^ strain, sharing the same genetic background, was used as a control.

Following the strategy used for the cellular models, we initially assessed autophagic markers. As shown in Fig. 7A, *dj-1β* KO flies exhibited higher levels of ubiquitylated proteins compared to controls, whereas, consistent with the cellular data, levels of Ref(2)p, the *Drosophila* ortholog of p62, remained unchanged. Accumulation of lipidated Atg8, ortholog of LC3, was also observed, in agreement with data from the cellular models (Fig. 7A). Given the relevance of DJ-1 to neurodegeneration, we repeated these analyses using head lysates, confirming that dj-1β deficiency also leads to the accumulation of ubiquitylated proteins and shows a trend toward higher Atg8-II levels in this tissue in comparison to control flies (Fig. 7B).

**Figure 7.**
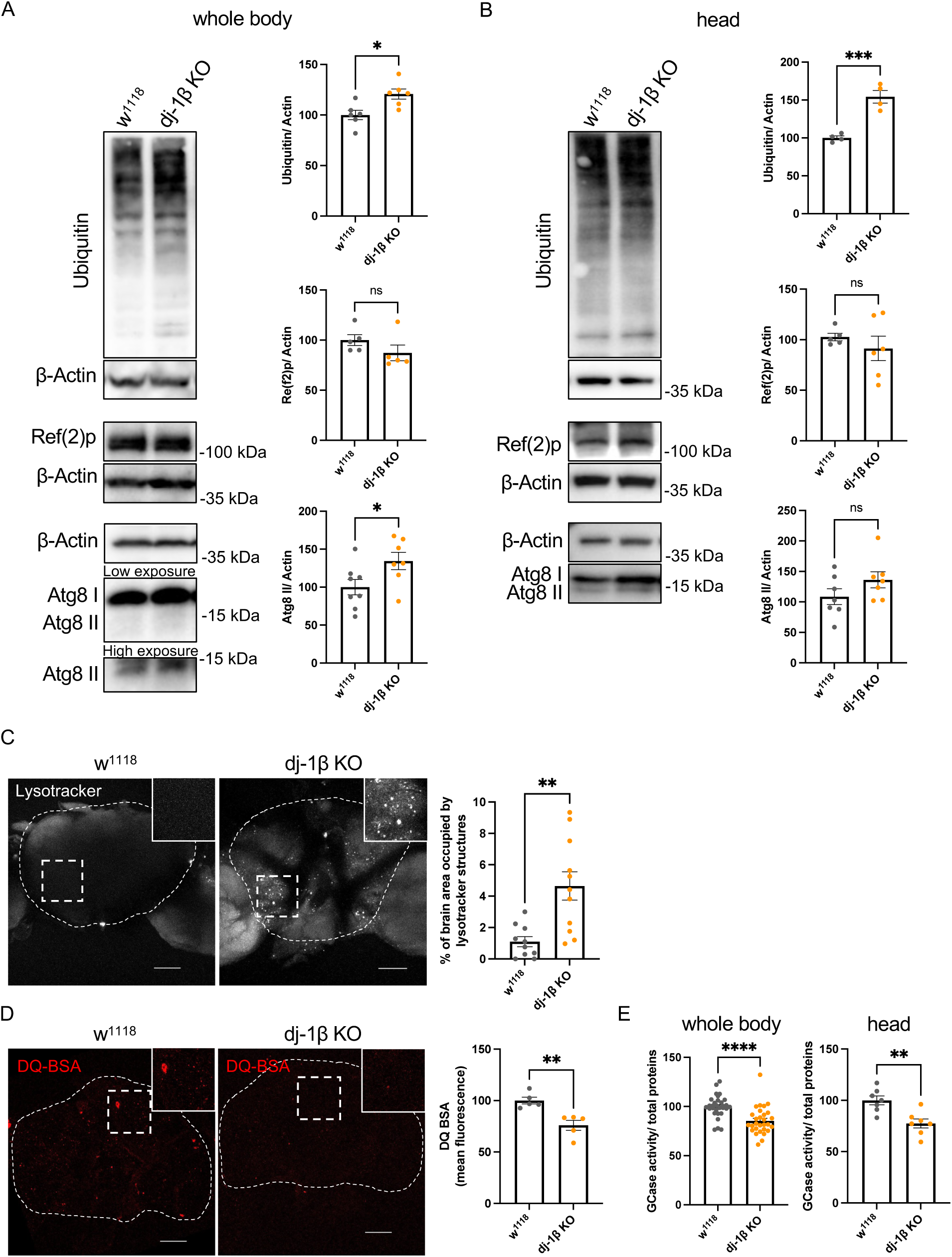
Loss of DJ-1 impairs autophagy and lysosomal function in fruit flies. (A) Western blot analysis of ubiquitinated proteins, Ref(2)p and Atg8 I/II in control (w^1118^) and dj-1β KO fly whole bodies. At least 5 independent biological replicates (pool of 5 flies per sample) were analyzed. Statistical significance was calculated using an unpaired t-test (Ubiquitin: t: 3.036; df: 10; * p < 0.05; Ref2p: t: 1.331; df: 8; not significant; lipidated Atg8 (Atg8 II): t: 2.275; df: 13; * p < 0.05). (B) Western blot analysis of ubiquitinated proteins, Ref(2)p and Atg8 in w^1118^ and dj-1β KO fly heads. At least 5 biological replicates (pool of 15 fly heads) were analyzed. Statistical significance was calculated using an unpaired t-test (Ubiquitin: t: 6.228; df: 6; *** p < 0.001; Ref(2)p: t: 0.8209; df: 9; not significant; Atg8 II: t:1.496; df: 12; not significant). (C) Confocal imaging of w^1118^ and dj-1β KO fly brains stained with LysoTracker-red. Scale bar: 50 μm. At least 10 brains per genotype were quantified. The panel on the right shows the quantification of the percentage of brain area occupied by LysoTracker-positive structures. Statistical significance was determined using an unpaired t-test (t: 3.557; df: 19; *** p < 0.001). (D) Confocal imaging of w^1118^ and dj-1β KO fly brains stained with the lysosomal proteases activity probe DQ-BSA. Scale bar: 50 μm. The mean DQ-BSA fluorescence intensity within the brains was analyzed in 5 brains per genotype. Statistical significance was determined using an unpaired t-test (t: 4.091; df: 8; ** p < 0.01). (E) GCase activity in w^1118^ and dj-1β KO fly whole bodies and heads. At least 30 biological samples per genotype for whole bodies (pools of 5 whole fly bodies per sample) and 7 biological replicates per genotype were analyzed for heads (pools of 15 heads per sample). Statistical analysis was performed using an unpaired t-test (whole body: t: 4.490: df: 58; **** p < 0.0001; heads: t:3.682; df: 12; ** p < 0.01).

We then examined lysosomal alterations in *dj-1β* KO flies. LysoTracker Red staining of fly brains revealed abnormal accumulation of enlarged acidic compartments in dj-1-deficient animals, as previously demonstrated in [31] and consistent with observations in MEFs and iPSCs (Fig. 7C). Moreover, DQ-BSA assays performed on fly brains demonstrated lower lysosomal protease activity in mutant flies relative to controls, indicating impaired lysosomal degradation due to dj-1β loss (Fig. 7D). To independently confirm lysosomal dysfunction, we also measured the activity of the lysosomal enzyme GCase. Given that heterozygous *GBA1* mutations are a known PD risk factor, GCase represents both a marker of lysosomal function and a PD-relevant parameter. Enzymatic assays in whole-body lysates showed significantly lower GCase activity in *dj-1β* null flies relative to control flies (Fig. 7E). Importantly, similar reductions were observed in lysates from *dj-1β* KO fly heads compared to controls (Fig. 7E).

### The absence of DJ-1 ortholog alters the AMPK and mTORC1 axis in fruit flies

To complete the *in vivo* validation of the cellular data, we assessed whether AMPK and/or mTORC1 signaling were altered by dj-1β loss in flies. In the absence of a suitable antibody to detect total *Drosophila* AMPK, we assessed AMPK activity by exclusively measuring its phosphorylation at Thr172. Analysis of whole-body protein lysates revealed a reduction in the active, phosphorylated form of AMPK in dj-1β KO flies compared with controls (Fig. 8A). To directly link this effect to dj-1β deficiency, we used the GAL4-UAS system [47] to ubiquitously overexpress dj-1β and subsequently measured p-AMPK levels. Notably, dj-1β overexpression led to a significant increase in p-AMPK compared with control flies (Fig. 8A), supporting a functional relationship between dj-1β levels and AMPK activation.

**Figure 8.**
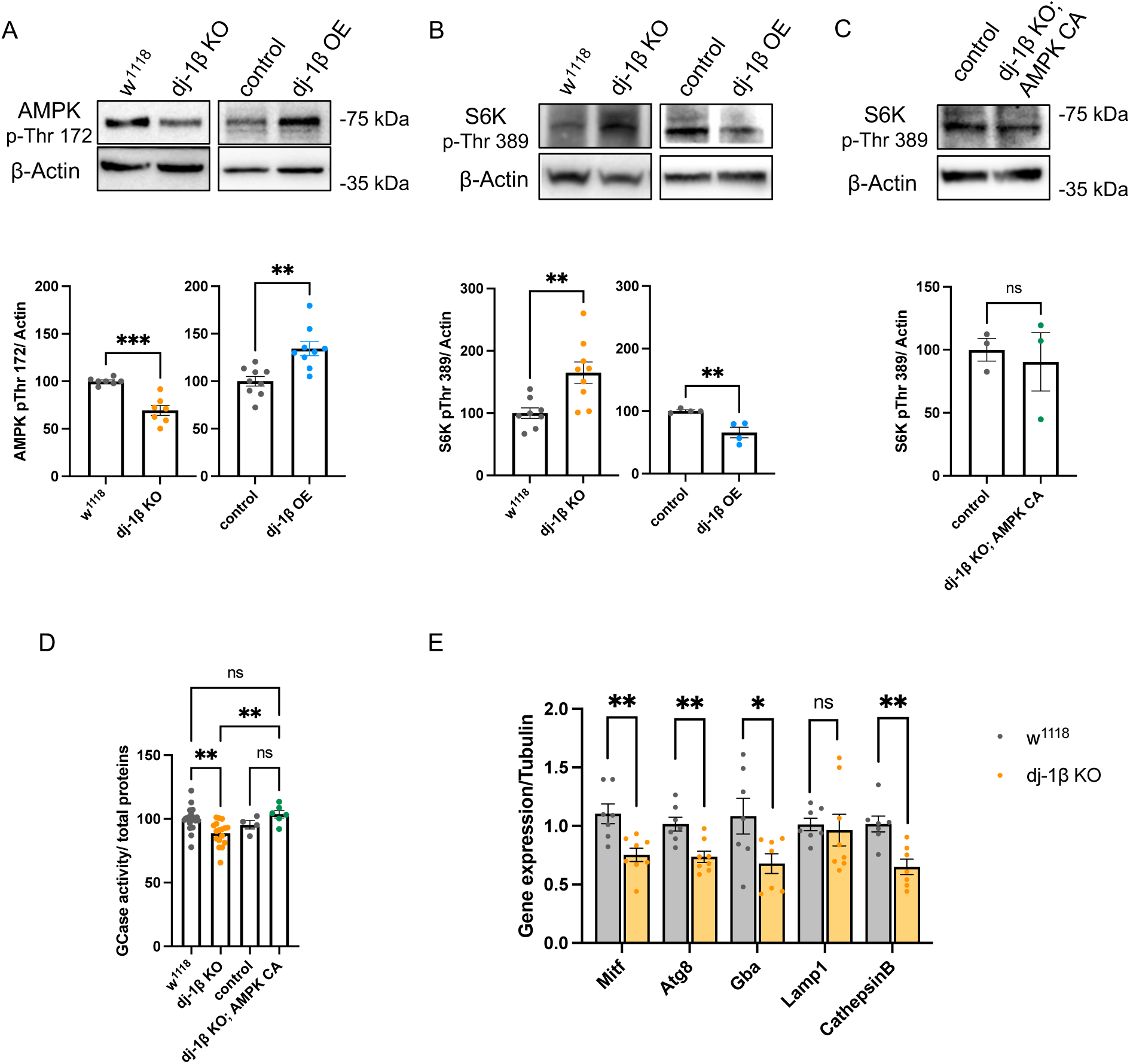
DJ-1 modulates the AMPK-mTORC1 pathway in flies. (A) Western blot analysis of phosphorylated AMPK (Thr172) in dj-1β KO, dj-1β overexpressing fly and genetically matched WT controls. The analysis was performed using at least 7 whole body samples per genotype (pools of 5 whole fly bodies per sample) Statistical significance was determined by an unpaired t-test (control vs dj-1β KO: t:5.692; df: 12; *** p < 0.001; control vs dj-1β OE: t: 3.851; df: 16; ** p < 0.01). (B) Western blot analysis of phosphorylated S6K (Thr389) in dj-1β KO, dj-1β overexpressing fly and genetically matched WT controls. The analysis was performed using at least 4 whole body samples per genotype (pools of 5 whole fly bodies per sample) Statistical significance was determined by an unpaired t-test (control vs dj-1β KO: t:5.692; df: 12; *** p < 0.001; control vs dj-1β OE: t: 3.851; df: 16; ** p < 0.01). (C) Western blot analysis of phosphorylated S6K (Thr172) in flies dj-1β KO overexpressing constitutive active AMPK (AMPK CA) and genetically matched WT control flies. The analysis was performed on 3 independent whole-body samples per genotype (pools of 5 whole fly bodies per sample) Statistical significance was determined by an unpaired t-test (t: 0.3863; df: 4; not significant). (D) GCase activity in dj-1β KO flies, dj-1β KO flies overexpressing AMPK CA and genetically matched WT control fly whole bodies. Statistical analysis was performed using a one-way ANOVA with Tukey’s multiple comparisons test (w^1118^ vs dj-1β KO: df: 42; ** p < 0.01; control vs dj-1β KO AMPK CA: df:42; not significant; w^1118^ vs dj-1β KO AMPK CA: df: 42; not significant; dj-1β KO vs dj-1β KO AMPK CA: df: 42; ** p < 0.01). (E) Real-time PCR analysis of genes regulated by TFEB in control and dj-1β KO fly heads. Gene levels were normalized to the housekeeping gene Tubulin. At least 7 independent biological replicates (pool of 15 fly heads) were analyzed. For each gene, the statistical significance between w^1118^ and dj-1β KO fly heads were determined using an unpaired t-test (Mitf: t: 3.573; df: 13; ** p < 0.01; Atg8: t: 3.729; df: 13; ** p < 0.01; Gba: t: 2.314; df: 12; *p < 0.05; Lamp1: t: 0.3112; df: 13; not significant; Cathepsin B: t: 3.888; df: 12; ** p < 0.01).

We next investigated the mTORC1 pathway. Owing to the lack of antibodies recognizing the non-phosphorylated form of the *Drosophila* S6K ortholog, we evaluated mTORC1 activity by measuring exclusively p-S6K levels. Consistent with our cellular data, WB analysis showed elevated p-S6K levels in dj-1β KO flies relative to controls, indicating enhanced mTORC1 activity upon dj-1β loss (Fig. 8B). Ubiquitous dj-1β overexpression reversed this effect, restoring p-S6K levels toward those observed in control flies.

To determine whether AMPK modulates mTORC1 signaling in this context, we generated flies expressing a constitutively active form of AMPK (AMPK-CA) [48] in the dj-1β KO background. Taking advantage of this mutant fly line, we observed the restoration of mTORC1 activity to the levels of control flies upon AMPK-CA overexpression (Fig. 8C). Importantly, AMPK-CA expression also rescued the reduction in GCase activity observed in KO flies (Fig. 8D).

To substantiate the functional impairment of the mTORC1 pathway *in vivo*, we then examined the transcriptional activity of Mitf, the *Drosophila* ortholog of TFEB. Real-time PCR analyses revealed that several Mitf-regulated genes, including *Mitf* itself, were downregulated in dj-1β –deficient flies, mirroring observations in MEFs (Fig. 8E).

### DJ-1-dependent redox alterations drive AMPK-mTORC1 pathway impairment both in MEF cells and fruit flies

The data collected so far suggest that DJ-1 participates in the regulation of the ALP by affecting both AMPK and mTORC1. Given the demonstrated role of DJ-1 in protecting against oxidative stress [49], and since ROS levels represent a factor able to affect AMPK activation [39,40], we next assessed whether altered redox conditions contributed to the DJ-1-dependent AMPK/mTORC1 dysregulation described here. Using the fluorescent dye dihydroethidium (DHE), which detects superoxide anions and hydrogen peroxide, we confirmed higher ROS levels both in *DJ-1* KO MEF and *dj-1β* KO fly brains relative to the corresponding controls, an effect that, in both cases, was rescued by treatment with the antioxidant N-acetylcysteine (NAC) (Fig. 9A-B) [31]. Importantly, ROS scavenging via NAC treatment was sufficient to restore AMPK phosphorylation and mTORC1 activity to levels comparable to those of the control in MEF cells (Fig. 9C) and in flies (Fig. 9D). These results not only confirm that the DJ-1 dependent increase of ROS represents one of the triggering factors modulating autophagy in DJ-1–deficient models, but also show that elevated ROS levels downregulate AMPK activity and promote mTORC1 signaling.

**Figure 9.**
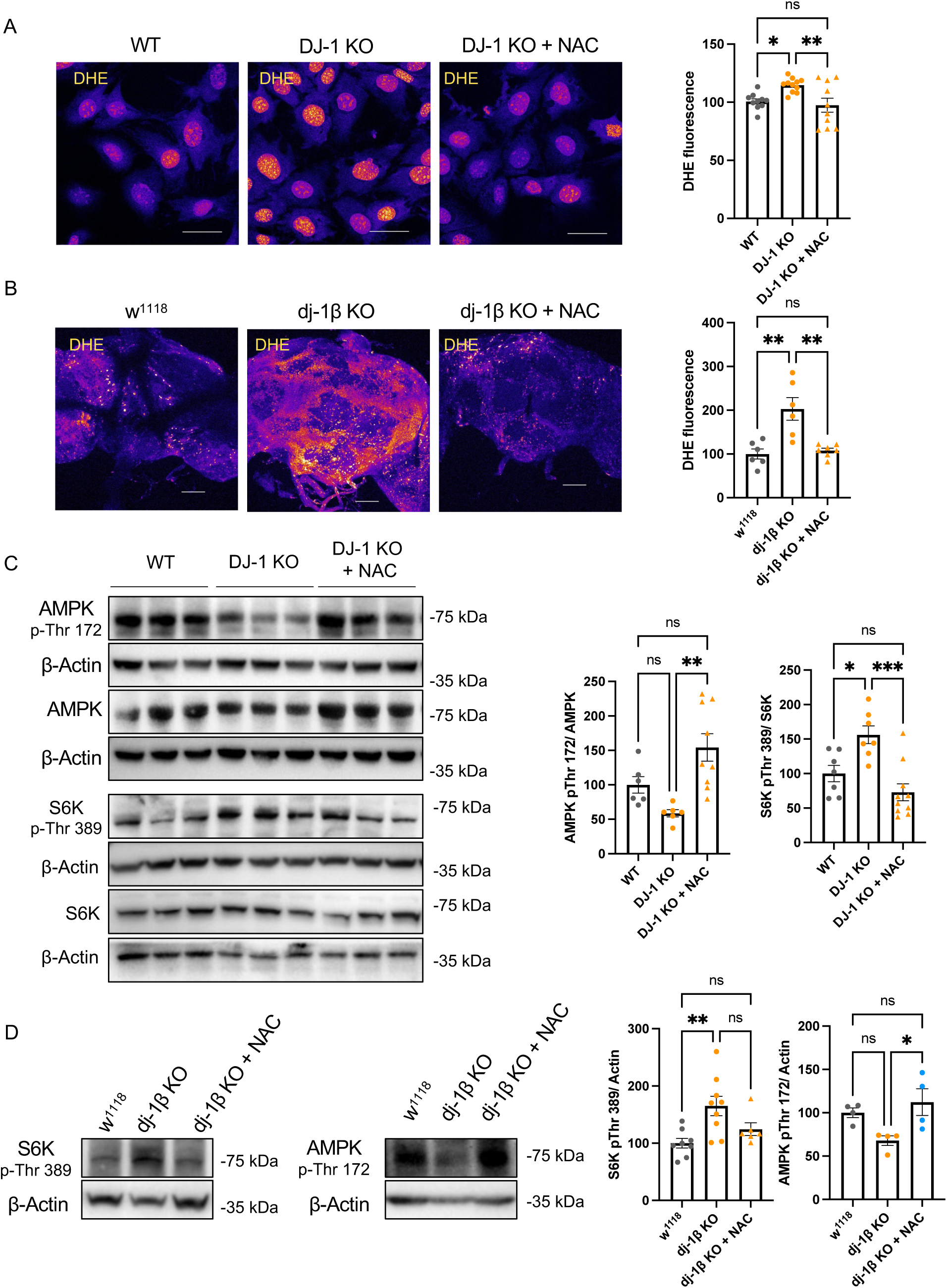
DJ-1-dependent ROS increase modulates the AMPK-mTORC1 pathway. (C) Live confocal imaging of MEF cells WT, DJ-1 KO and DJ-1 KO treated with NAC stained with the ROS reporter DHE. Scale bar: 50 μm. The mean DHE fluorescence intensity was quantified in every confocal field. The analysis includes at least 10 fields of 3 independent batches of cells. Statistical significance was determined using a one-way ANOVA with Tukey’s multiple comparisons test (WT vs DJ-1 KO: df: 28 * p < 0.05; WT vs DJ-1 KO + NAC: df: 28; ** p < 0.01; DJ-1 vs DJ-1 KO + NAC; df: 28; not significant) (B) Confocal imaging of w^1118^, dj-1β KO and dj-1β KO flies treated with NAC and stained with the ROS reporter DHE. Scale bar: 50 μm. At least 6 brains per condition were included in the analysis. Quantification of DHE fluorescence integrated density is shown in the graph on the right. Statistical significance was performed using one-way ANOVA with Tukey’s multiple comparisons test (w^1118^ vs dj-1β KO: df: 16; ** p < 0.01; w^1118^ vs dj-1β KO + NAC: df: 16; not significant; dj-1β KO vs dj-1β KO + NAC: df: 16; ** p < 0.01). (C) Western blot analysis of total and phosphorylated AMPK (Thr172) and total and phosphorylated S6K (Thr389) in WT, DJ-1 KO and DJ-1 KO MEF cells treated with the antioxidant NAC (1 mM, 24 h). At least 6 independent samples were analyzed for AMPK and 7 independent samples for S6K evaluation. The graphs on the right show the level of AMPK phosphorylation normalized to total AMPK (the analysis was performed using one-way ANOVA with Tukey’s multiple comparisons test. WT vs DJ-1 KO: df: 18; not significant; WT vs DJ-1 KO + NAC: df: 18; not significant; DJ-1 KO vs DJ-1 KO + NAC; df: 18; ** p < 0.01) and the phosphorylation of S6K normalized to total S6K level (one-way ANOVA with Tukey’s multiple comparisons test. WT vs DJ-1 KO: df: 21; *p < 0.05; WT vs DJ-1 KO + NAC: df: 21; not significant; DJ-1 KO vs DJ-1 KO + NAC; df: 21; *** p < 0.001). (D) Western blot analysis of phosphorylated AMPK and phosphorylated S6K (Thr389, mTORC1 target site) in w^1118^, dj-1β KO and dj-1β KO flies treated with NAC (10 mM, 24h). At least 6 samples were analyzed (pools of 5 whole bodies). The graphs show the level of phosphorylated AMPK (Statistical significance was determined using one-way ANOVA with Tukey’s multiple comparisons test. w^1118^ vs dj-1β KO: df: 9; not significant; w^1118^ vs dj-1β KO + NAC: df: 9; not significant; dj-1β KO vs dj-1β KO + NAC: df: 9; * p < 0.05) and phosphorylated S6K (one-way ANOVA with Tukey’s multiple comparisons test. w^1118^ vs dj-1β KO: df: 20; ** p < 0.01; control vs dj-1β KO + NAC: df: 20; not significant; dj-1β KO vs dj-1β KO + NAC: df: 20; not significant).

## Discussion

The PD-associated protein DJ-1 has long been described to protect cells against excessive ROS levels [50,51], a function possibly supported by the participation of the protein in the maintenance of mitochondrial homeostasis. Indeed, under oxidative conditions, DJ-1 has been shown to translocate inside mitochondria [52–54], where it appears to preserve the organelle functionality, by interacting and sustaining the activity of complex I [15,16,31] and complex V [17]. Additionally, the protein has been found to take part in the process of mitochondrial quality control, playing a role in the Parkin/PINK1-mediated mitophagy [14,19,21,23,24], and in the regulation of the mitochondrial dynamics [13,18,20,22]. While the participation of DJ-1 in the maintenance of mitochondrial homeostasis is well accepted, its involvement in autophagy is much less characterized.

The data presented here allow us to propose a mechanism that links DJ-1 function to the regulation of the autophagy-lysosomal pathway, providing insights with direct relevance to the pathology of PD. By leveraging three distinct experimental systems, we have elucidated a signaling cascade that describes an intracellular communication pathway between mitochondria and lysosomes.

Our findings in all DJ-1-deficient models consistently pointed to the accumulation of substrates targeted for degradation, a primary indicator of a breakdown in cellular quality control. More specifically, higher levels of ubiquitylated proteins and lipidated LC3 were detected, even though we did not observe variations in p62 protein levels. It is worth mentioning here that although p62 is frequently used as a reporter of autophagy, its levels are tightly regulated at both transcriptional and post-transcriptional levels by multiple cellular pathways; therefore, its abundance does not always directly reflect autophagic flux [55]. Accordingly, we observed that p62 gene transcription levels are significantly reduced in DJ-1 KO cells. The accumulation of autophagic markers was accompanied, in each experimental model, by increased lysosomal mass. Microscopically, this manifested as an abnormal accumulation of acidic compartments in the brains of *dj-1* null flies and a higher number of LAMP1-positive structures in MEFs and dopaminergic neurons. Crucially, this increase in lysosomal mass was associated with a sharp decrease in lysosomal proteolytic activity. Thus, our data suggest a critical dichotomy between the size of the lysosomal compartment and lysosomal functionality, where DJ-1 deficiency leads to the accumulation of enlarged but enzymatically hypoactive lysosomes. This was corroborated by transmission electron microscopy, which revealed the accumulation of structures containing electron-dense material, characteristic of undigested cellular cargoes, that were absent in control cells.

Mechanistically, our investigation revealed that the loss of DJ-1 results in persistent and aberrant inhibition of AMPK, leading to consequent activation of mTORC1. Hyperactivation of mTORC1 exerted detrimental effects on key downstream autophagic regulators: it directly inhibited the autophagy-initiating kinase ULK1 through phosphorylation and promoted both degradation and cytoplasmic retention of TFEB (and its *Drosophila* ortholog Mitf), the master transcriptional regulator of autophagy and lysosomal biogenesis. Consistently, DJ-1 KO MEFs displayed a marked reduction in total TFEB protein levels together with impaired nuclear translocation of the remaining TFEB. In agreement with these findings, real-time PCR analysis revealed significant downregulation of TFEB/Mitf target genes in both MEFs and flies. importantly, the rescue of mTORC1 hyperactivation upon expression of constitutively active AMPK in dj-1 null flies indicates that AMPK acts upstream of mTORC1.

The role of AMPK as a key mediator of long-distance communication between mitochondria and the ALP has been previously described [11]. AMPK is primarily regulated by ATP levels [12], making it a central sensor of mitochondrial energy generation. In turn, it promotes autophagy through modulation of both mTORC1 and TFEB. Notably, we previously demonstrated that ATP levels are not altered in our DJ-1 knockout *Drosophila* models, suggesting that AMPK inhibition in the absence of DJ-1 is driven by alternative stimuli [31].

In this context, our experiments establish a mechanistic link between the known role of DJ-1 in redox control and the autophagy defects observed in its absence. Consistent with previous findings [31], DHE staining showed that deletion of dj-1 induces significant oxidative stress, characterized by increased ROS levels in both cellular and fly models. Notably, pharmacological intervention with the antioxidant compound NAC demonstrated a causal relationship: treatment of DJ-1 KO cells and dj-1 null flies with NAC not only normalized ROS levels but also fully restored AMPK phosphorylation levels and the activity of its downstream target mTORC1.

The relationship between ROS and AMPK activity remains an area of active investigation [39]. While several studies report that elevated ROS levels promote AMPK activation [56,57], others have described the opposite effect [58]. Our observation of an inverse correlation between ROS levels and AMPK phosphorylation is therefore not necessarily inconsistent with previous findings. Rather, we propose that both the magnitude and the duration of ROS signaling may differentially influence AMPK activity and its downstream pathways.

The relatively modest oxidative stress induced by DJ-1 loss in our models is consistent with previous studies linking redox imbalance to the regulation of autophagy. For example, low ROS levels, similar to those observed under DJ-1 KO conditions, have been reported to stimulate mTORC1 activity, whereas high ROS concentrations or prolonged oxidative stress inhibit it [59]. Likewise, studies of mitochondrial respiratory chain deficiency have shown that AMPK activity can vary depending on the nature of the mitochondrial insult: reduced AMPK activation has been observed during chronic mitochondrial dysfunction, whereas increased AMPK activity occurs in response to acute mitochondrial impairment [11].

While a ROS-independent role of DJ-1 in mitophagy was recently described in the presence of valinomycin, a mitochondrial uncoupler that promotes mitophagy [24], a notable aspect of this study is that the results were consistently obtained under basal conditions. As a consequence, although the magnitude of the measured effects was modest, unsurprisingly given that DJ-1 is not a vital protein, they nonetheless highlight pathways modulated by the protein. Even though such alterations might be easily tolerated under normal conditions, they may become detrimental when sustained over time, especially in post-mitotic neuronal cells.

In conclusion, this research identifies DJ-1 as a critical homeostatic regulator that maintains the integrity of the autophagy-lysosomal pathway by preventing redox-imbalance-driven AMPK inhibition and mTORC1 hyperactivation and provides a compelling molecular explanation for the dual mitochondrial/lysosomal defects that are known hallmarks of DJ-1-associated Parkinsonism.

## Materials and Methods

### MEF cell cultures

Mouse embryonic fibroblast (MEF) cells are a generous gift from Prof. Marisa Brini’s lab, that generated them from WT and DJ-1 KO mice used in [60]. MEF cells were cultured in Dulbecco’s modified. Eagle’s medium (DMEM, Life Technologies), supplemented with 10% fetal bovine serum (FBS, Life Technologies). Cell lines were maintained at 37°C in a 5% CO2 controlled atmosphere. TrypLE Express Enzyme (Life Technologies) was employed to generate subcultures. The absence of DJ-1 was confirmed through western blot. Mycoplasma testing was regularly performed using mycoplasma PCR kit (Euroclone).

### iPS cell culture and differentiation in dopaminergic neurons

DJ-1 KO induced pluripotent stem cells (iPSC) were generated in [44] from the Human Episomal control iPSC line (Gibco, A18945). The absence of DJ-1 was confirmed via western blot analysis. Cells were grown on plates pre-coated with Matrigel (CORNING 356231) in E8 media (Thermo Scientific, cat #A1517001). 10uM Rock inhibitor (STEMCELL, Cat # 72304) was added to the media for about 24 h after splitting and thawing. Media was changed every day. TrypLE Express Enzyme (Life Technologies) was used to detach cells and generate subculture. Differentiation into dopaminergic neurons was carried out following the protocol explained in https://dx.doi.org/10.17504/protocols.io.bsq5ndy6 until day 25. Differentiation was confirmed via western blotting analysis of the dopaminergic neuronal marker tyrosine hydroxylases.

### *Drosophila* strains and husbandry

*Drosophila* w^1118^ (BDSC_5905), dj-1βΔ93 (BDSC_33601), UAS-dj-1β (BDSC_33604), UAS-AMPK T184D (BDSC_32110), daughterless (da)-Gal4 (BDSC_8641) fly lines were obtained from the Bloomington *Drosophila* Stock Center. All strains were reared on common cornmeal food in a humidified, temperature-controlled incubator at 25 °C on a 12 h light/dark cycle. For each experiment, 1-to-3-day-old male flies were used.

### Drosophila treatments

1–3-day old adult fly males were collected under brief CO2 exposure and transferred into new tubes containing standard food (20 flies/vial) supplemented 10 mM of NAC (Sigma Aldrich: A7250) for 24 hours.

### Cell treatments

MEF cells were treated with 1 mM of NAC (Sigma Aldrich: A7250) for 24 hours.

### Cells transfection

MEF cells were transfected with plasmid DNA of WT-DJ-1 and mCherry-LC3-GFP reporter using Lipofectamine 2000 (Invitrogen) for 48 hours, following the manufacturer’s instructions.

### Cell lysis and Western blotting

Fly tissues or cells were lysed in RIPA buffer (20 mM Tris-HCl pH 7.5, 150 mM NaCl, 1 mM EDTA, 1 mM sodium orthovanadate, 1 mM β-glycerophosphate, 2,5 mM sodium pyrophosphate, 1% Tween) supplemented with protease inhibitors (Roche) and incubated on ice for 30 minutes. Lysates were cleared by centrifugation at 20000 x g for 30 minutes at 4°C. Supernatants were used. For each sample, 30-50 μg of protein were loaded on or ExpressPlus™ PAGE 4–20% gels (GenScript), in MOPS running buffer or 15% Tris-glycine polyacrylamide gels in SDS/Tris-glycine running buffer. The resolved proteins were transferred to polyvinylidenedifluoride (PVDF) membranes (Bio-Rad) with the Trans-Blot® Turbo™ Transfer System (Bio-Rad). Membranes were blocked in Tris-buffered saline plus 0.1% Tween (TBS-T) plus 5% bovine serum albumine (BSA-Sigma aldrich) 1 h at room temperature (RT) and then incubated over-night at 4 °C with primary antibodies diluted in TBS-T plus 5% BSA. Membranes were then incubated for 1h at RT with horseradish peroxidase (HRP)-conjugated secondary antibodies. Immunoreactive proteins were visualized using Immobilon® Forte Western HRP Substrate (Merck Millipore) at the Imager Chemi Premium (VWR). Densitometric analysis was carried out using the Image J software.

Antibodies utilized for western blot:

fly samples: mouse β-actin (Sigma Prestige); rabbit Ref(2)p (Abcam: ab178440); mouse Ubiquitin (Santa Cruz Biotech: sc-8017), rabbit p-S6K (Thr389)(Cell Signaling technology: 9205), rabbit GABARAP/Atg8a (Abcam: ab109364).

iPSC: rabbit p62 (Abcam: ab109012), rabbit LC3 (Abcam: ab192890), mouse LAMP1 (Abcam: ab25630), rabbit TH (Pel-freez biological: P40101-150), rabbit S6K (Cell Signaling technology: 9202), rabbit p-S6K (Thr389) (Cell Signaling technology: 9205), rabbit DJ-1 (Abcam: ab18257).

MEF cells: mouse β-actin (Sigma-Aldrich, A1978), rabbit PARP (Cell Signalling Technology, 9542S), rabbit TFEB (Bethyl Laboratories, A303-673A), mouse GAPDH (Sigma-Aldrich: G8795) rabbit LC3 (Abcam: ab192890), mouse Ubiquitin (Santa Cruz Biotech: sc-8017), rat LAMP1 (XXX), guinea pig p62 (Progen: GP62-C), rabbit S6K (Cell Signaling technology: 9202), rabbit p-S6K (Thr389)(Cell Signaling technology: 9205), rabbit ULK1 (Cell Signaling technology: 8054), rabbit p-ULK1 (Ser757)(Cell Signaling technology: 6888), rabbit DJ-1 (Abcam: ab18257), rat LAMP1 (Abcam: ab25245).

### Cell fractionation

Fractionation of cell nuclear and cytoplasmic compartments was performed using the NE-PERTM Nuclear and Cytoplasmic Extraction Reagents (Thermo-Scientific) kit according to the manufacturer’s instructions. Samples were analysed via western blotting as previously explained.

### TEM imaging

Cells were plated onto 6 well-plates and cultured until they reached about 80% of confluency. Cells were then fixed for 20 minutes in 0.1 M sodium cacodylate (pH 7.4), containing 2.5% glutaraldehyde and 2% paraformaldehyde, and then dissected to isolate thoraces. Briefly, the head, legs, and wings were. Samples were incubated with a solution of 1% of tannic acid for 1 h at room temperature and then post-fixed with 1% osmium tetroxide in 0.1 M sodium cacodylate buffer for 1h at 4 °C. After three water washes, samples were dehydrated in a graded ethanol series and embedded in an epoxy resin (Sigma-Aldrich). Ultrathin sections (60–70 nm) were obtained with an Ultrotome V (LKB) ultramicrotome, counterstained with uranyl acetate and lead citrate and viewed using a Tecnai G2 (FEI) transmission electron microscope, operating at 100 kV. Images were captured with a Veleta (Olympus Soft Imaging System) digital camera.

### Immunocytochemistry and confocal imaging

Cells were cultured onto 12 mm glass coverslips (Thermo-Scientific) coated with 10 μg/ml Laminin (Sigma Aldrich: L2020) and 2 μg/ml Fibronectin (Corning: 356008) for iPSC-derived neurons and with Poly-L-Lysine (Sigma-Aldrich) for MEF cells. Cells were washed in PBS and fixed with 4% w/v Paraformaldehyde (PFA) for 20 minutes at room temperature (RT). After cell permeabilization in PBS plus triton 0,1% for 20 minutes at RT and a blocking step performed in 3% BSA diluted in PBS for 30 minutes at RT, cells were stained with the appropriate primary antibody diluted in PBS plus 3% BSA overnight at 4°C. Subsequently, cells incubated with the secondary antibody Alexa Fluor® 488 conjugated (Life Technologies), Alexa Fluor® 568 conjugated (Life Technologies) or Alexa Fluor® 647 for 1 hour at RT. Before mounting the coverslips on glass slides, cells were incubated with Hoechst 33258 (Invitrogen) diluted in PBS for 5 minutes. Images were acquired with a LeicaSP8 confocal microscope (Leica Microsystems) and quantified using ImageJ. The following primary antibodies were used:

iPSC: sheep TH (Pel-freez biological: P60101-150), mouse LAMP1 (Abcam: ab25630), rabbit LC3 (Abcam: ab192890), Ubiquitin (Santa Cruz Biotech: sc-8017).

MEF cells: rat LAMP1 (Abcam: ab25245).

### Lysotracker staining

MEF cells: Cells were culture onto 6-wells plates. When they reached 90% of confluency cells were treated with 100 nM LysoTracker™ Green DND-99 (Thermofisher Scientific, L7526) for 1 hour. Cells were then harvested in PBS and analysed by flow cytometry.

*Drosophila*: adult male brains were dissected in phosphate-buffered saline (PBS) followed by 15 minutes of incubation with 200 nM LysoTracker™ Red DND-99 probe (Thermofisher Scientific, L7528) at RT in the dark. Subsequently, brains were washed trice in PBS for 5 minutes, mounted in Mowiol ® 4-88 (Calbiochem, 475904), and immediately analyzed. Z-stack images were acquired on a LeicaSP5 confocal microscope (Leica Microsystems). The area of the brain occupied by LysoTracker staining and the number of acidic compartments were quantified by ImageJ.

### DQ-BSA assay

MEF cells: cells: Cells were culture onto 6-wells plates. When they reached 90% of confluency cells were treated with 10 μg/ml DQ-BSA probe (Thermofisher: D12051) for 45 minutes in the dark at 37 C° followed by 2 hours of chasing. Cells were then harvested in PBS and analysed by flow cytometry.

iPSC-derived neurons: cells were cultured onto an 8 well-chambered coverslip for cell culture and live imaging (ibidi: 80806) coated with 10 μg/ml Laminin (Sigma Aldrich: L2020) and 2 μg/ml Fibronectin (Corning: 356008). Cells were incubated with 10 μg/ml of DQ-BSA probe (Thermofisher: D12051) diluted in PBS for 45 minutes in the dark at 37 C°. Cells were then chased for 2 hours in DMEM. Finally, cells were incubated in DMEM without phenol red (Life Technologies) and observed live with a Zeiss LSM 700 confocal microscope. DQ-BSA probe fluorescence intensity was quantified with ImageJ.

*Drosophila*: 1-3 days old male fly brains were dissected in Schneider’s Drosophila medium (Biowest, L0207) and incubated with 80 μg/ml, for 60 minutes at RT in the dark. Brains were then rinsed 3 times and chased for 2 hours at RT in Schneider’s Drosophila medium before being mounted in Mowiol® 4-88 (Calbiochem, 475,904). Images were acquired with LeicaSP5 confocal microscope (Leica Microsystems) and quantified using ImageJ.

### Cytofluorimeter analysis

Flow cytometric analysis was performed using the BD LSRFortessa™ X-20 Cell Analyzer (BD Biosciences). Lysotracker Green and DQ-BSA Red signals were detected in the FITC and APC channels, respectively.

Untreated cells were used as negative controls, while cells stained individually with either Lysotracker or DQ-BSA were used to set detector voltages and define acquisition parameters.

### DHE assay

MEF cells: Cells were cultured onto 8 wells-plates for live imaging (Ibidi). Cells were incubated with 30 μM dihydroethidium (DHE) probe (Invitrogen, D11347) for 1 hour in the dark at 37 C°. Subsequently cells were washed in PBS and then incubated in DMEM without phenol red (Life Technologies) for live imaging visualization. Images were acquired utilizing LeicaSP8 confocal microscope (Leica Microsystems) and analysed with ImageJ.

*Drosophila:* adult male brains were dissected in Schneider’s Drosophila medium (Biowest, L0207) followed by 5 min of incubation with 30 μM dihydroethidium (DHE) probe (Invitrogen, D11347) in the dark. Subsequently, brains were washed thrice in Schneider’s Drosophila medium for 5 min and mounted in Mowiol ® 4-88 (Calbiochem, 475904). Images were acquired utilizing LeicaSP5 confocal microscope (Leica Microsystems). The intensity of DHE staining was quantified by ImageJ.

### MagicRed staining

Cells were cultured onto 8 wells-plates for live imaging (Ibidi). When cells reached 90% of confluency were incubated with MagicRed Cathepsin B activity following manufacturer instruction. Briefly, cells were incubated with the probe for 1 hour in the dark at 37 C°. Subsequently cells were washed in PBS and then incubated in DMEM without phenol red (Life Technologies) for live imaging visualization. Images were acquired utilizing LeicaSP8 confocal microscope (Leica Microsystems) and analysed with ImageJ.

### mCherry-GFP-LC3 staining and live imaging

Cells were cultured onto 8 wells-plates for live imaging (Ibidi). When cells reached 50 % of confluency were transfected with the mCherry-GFP-LC3 plasmid using Lipofectamine 3000 (Invitrogen) according to manufacturer instruction. 36-48 hours post transfection cells were washed in PBS and then incubated in DMEM without phenol red (Life Technologies) for live imaging visualization. Images were acquired utilizing LeicaSP8 confocal microscope (Leica Microsystems) and analysed with ImageJ.

### Real time PCR

MEF cells: RNA was extracted from MEF cells using the RNeasy Mini Kit (Qiagen) following manufacturer’s instructions. RNA quality and concentration were assessed by Nanodrop. Complementary DNA (cDNA) was synthesized from total RNA using iScript cDNA Synthesis Kit (Bio Rad). qPCR reactions were performed on 98-well plate adding to the cDNA the PowerUp SYBR Green Master Mix (Applied Biosystems) and the DNA primers.

*Drosophila*: RNA was extracted from fly whole body or head tissues using the miRNeasy Mini Kit (Thermofisher: 217004) following manufacturer instuctions. RNA quality and concentration were assessed by Nanodrop. cDNA was synthesized from total RNA using 5 iScript cDNA Synthesis Kit (Bio Rad). qPCR reactions were performed on 98-well plate adding to the cDNA the PowerUp SYBR Green Master Mix (Applied Biosystems) and the DNA primers.

## Statistical analysis

Graphs and statistical analyses were performed by using GraphPad Prism 10. Generally, data are represented as scatter plots with bars showing mean with standard error of the mean (SEM). Statistical significance was defined for p-value <0.05 (ns p > 0.05, * p < 0.05, ** p < 0.01, *** p < 0.001, **** p < 0.0001). Additional details are reported in the legend of each figure.

## Acknowledgments

MB was supported by a grant from PR Veneto FESR 2021-2027 (RIxAA). FA was supported by a Boehringer Ingelheim travel grant.

## Abbreviation list

ALP: autophagic-lysosomal pathway
AMPK: AMP-activated protein kinase
AMPK-CA: constitutive active form of AMP-activated protein kinase
pAMPK: phosphorylated form of AMP-activated protein kinase
DHE: dihydroethidium
*GBA*1: glucocerebrosidase 1 gene
GCase: glucocerebrosidase 1 protein
KO: knockout
iPSCs: induced pluripotent stem cells
LC3: microtubule-associated protein 1A/1B-light chain
mTORC1: mechanistic target of rapamycin complex 1
NAC: N-acetylcysteine
PD: Parkinson disease
p-S6K: phosphorylated form of ribosomal protein S6 kinase beta-1
ROS: reactive oxygen species
S6K: ribosomal protein S6 kinase beta-1
TH: tyrosine hydroxylase
WB: western blot
WT: wild-type

